# Combining Deep Learning and Active Contours Opens The Way to Robust, Automated Analysis of Brain Cytoarchitectonics

**DOI:** 10.1101/297689

**Authors:** Konstantin Thierbach, Pierre-Louis Bazin, Walter De Back, Filippos Gavriilidis, Evgeniya Kirilina, Carsten Jäger, Markus Morawski, Stefan Geyer, Nikolaus Weiskopf, Nico Scherf

## Abstract

Deep learning has thoroughly changed the field of image analysis yielding impressive results whenever enough annotated data can be gathered. While partial annotation can be very fast, manual segmentation of 3D biological structures is tedious and error-prone. Additionally, high-level shape concepts such as topology or boundary smoothness are hard if not impossible to encode in Feedforward Neural Networks. Here we present a modular strategy for the accurate segmentation of neural cell bodies from light-sheet microscopy combining mixed-scale convolutional neural networks and topology-preserving geometric deformable models. We show that the network can be trained efficiently from simple cell centroid annotations, and that the final segmentation provides accurate cell detection and smooth segmentations that do not introduce further cell splitting or merging.

## 1 Introduction

Systematic studies of the cortical cytoarchitecture are indispensable to understand the functional organization of the human brain. Classical works based on qualitative description of cell counts and shapes in physical 2D sections of the human cortex revealed functional areas and segregation in the brain [2, 4, 15]. These brain parcellations are currently updated and refined using automated image analysis [18]. Even 3D imaging of post mortem brain tissue at microstructural resolution are within reach using recent light sheet fluorescence microscopy (LSFM) [7] and tissue clearing protocols [3]. Combined with advanced image analysis these techniques enable studying cortical cellular organisation in the human brain with unsurpassed precision. To reach this goal we need robust computational analysis relying on minimal manual annotations, facing the following challenges:

- Clearing of aged, unperfused human tissue is imperfect, and optical distortions due to scattering and refraction remain. This leads to varying background intensities across the image and shading artifacts.
- The penetration of antibody stains and thus contrast is uneven across the sample. The tissue degenerates with longer post-mortem times. These effects increase the already high variability of neural shape and appearance across the cortical samples (Fig. 1a).
- The resolution is lower along the optical axis in the 3D stack. Additional imperfection in depth focusing and sample movement create artifacts through the depth of the stack (Fig. 1b).
- Cell density varies locally, leading to false segmentation of cells into clusters.

Machine Learning methods improved the analysis of microscopy data [13]. Deep Learning, in particular Convolutional Neural Networks (CNNs), can address challenging problems in biomedical imaging because they learn multi-level internal representations of the data [9, 13]. These, typically supervised, methods require a lot of annotated data: For cell segmentation pixel-accurate masks have to be supplied [12]. Manually annotating data for training is often prohibitive in biomedical applications where data are specialized, scarce and expert knowledge is required. Abstract concepts at the object level (Gestalt principles such as continuation, closure [8], or object topology) are hard to learn with CNNs. Additional annotation of the border region between adjacent cells is needed to reduce false merging of neighboring cells [12]. Human vision exploits high level concepts using top-down processing [8] which is not represented in feedforward architectures.

Active Contour methods have been designed to embody high level concepts of object shapes. They can guarantee the smoothness of contours and a consistent topology [1]: features that improve cell segmentation in challenging conditions and prevent splitting and merging of contours during segmentation. But active contour methods require an initialization with the number and approximate position of objects in the image. Robust initial localization of cells is hard to define a priori and should be learned from data. This is where Deep Learning has a clear advantage: CNNs can be trained to robustly predict cell positions in images using only sparse centroid annotations [16].

In this work we combine the complementary strengths of CNNs and topology-aware active contours into a robust workflow to detect and segment cells that delivers high quality results and most importantly, requires only minimal annotations (sparse annotations of approximate cell centers are enough)^6^. Additionally, our approach works with 2D and 3D microscopy data. Here, we demonstrate and validate the method using 2D slices from a 3D microscopy image volume obtained of cleared post mortem human brain blocks and as a proof of concept we show that our methods easily extends to full 3D processing.

## 2 Methodology

### 2.1 Sample Preparation

Blocks from a human post mortem brain (temporal lobe cortex, male, 54 yr., post-mortem interval 96h) have been provided by the Brain Banking Centre Leipzig of the German Brain-Net. The entire procedure of case recruitment, acquisition of the patient’s personal data, the protocols and the informed consent forms, performing the autopsy, and handling the autoptic material have been approved by the local ethics committee. For details on tissue preparation and clearing see [10].

**Fig. 1.**
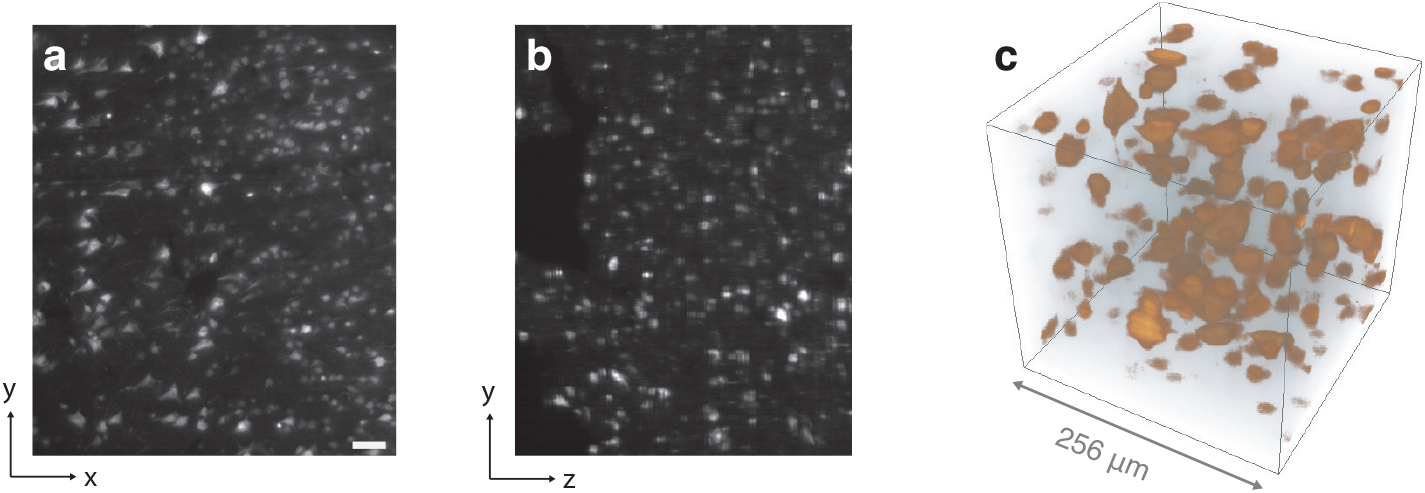
Example of image data. An xy (a) and yz (b) slice of a 1080 × 1280 × 1000*μ*m subvolume (scale bar 100*μ*m). (c) Direct volume rendering of cuboid volume randomly sampled from the image stack.

### 2.2 Image Data

A commercial light-sheet fluorescence microscope (LaVision BioTec, Bielefeld, Germany) was used to image the cleared specimen. The microscope was equipped with 10x CLARITY-objective (Olympus XLPLN10XSVMP, numerical aperture (NA) 0.6, working distance (WD) 8 mm; Zeiss Clr Plan-Apochromat, NA 0.5, WD 3.7 mm) and operated with 630 nm excitation wavelength and band-pass 680 nm emission filter. Samples were stained with a fluorescent monoclonal antibody against human neuronal protein HuC/HuD (a specific marker for neuronal cell bodies). The acquisition covered a 1.1 mm × 1.3 mm× 2.5 mm volume resulting in a stack of 2601 16-bit TIFF images (2560×2160 pixels, 0.51 *μ*m lateral resolution) using a 1 *μ*m step size.

For the 2D analysis workflow we took 19 slices at regular intervals from the entire stack. We used 15 images for training and validation and kept 4 images as a test set. The images for the test set were used for final assessment of segmentation and detection performance only. A single image typically contains around 300 cells (Fig. 1). To analyze 2D segmentation accuracy, an expert created reference cell masks on the 2D test images. The masks were independently checked and corrected by a second expert.

For the 3D analysis pipeline we first resampled the image stack to an isotropic resolution of 1 *μ*m. One expert manually annotated cell centroids for a subregion of the size 2304 × 256 × 1280 (z,x,y), which we subsequently used for the training and validation of the CNN for cell localization. We additionally annotated cell centroids in three separate regions of size 256 × 256 × 256, which served as a test set to measure the performance of our method. As manually segmenting cells in 3D is very laborious and error-prone we only annotated 2D reference cell masks in regularly spaced xy, and yz planes of the test images. To validate the agreement between annotated masks and segmentation we computed the segmentation entirely in 3D but exported the results only in those 2D planes that have been used for annotation.

### 2.3 Cell Segmentation Workflow

The proposed method is based on a Fully Convolutional Neural Network for cell localization and a topology-preserving multi-contour segmentation [1] to control smoothness and topology of the segmentation. To handle the different cell sizes we use the recently proposed Mixed-scale Dense (MS-D) architecture by [11] to robustly predict masks of cell centroid regions in 2D and 3D. The basic concept is as follows: **Training:** Pairs of image stacks (annotated centroids convolved by a spherical kernel and raw data) are fed into MS-D network. The network is trained to directly segment a spherical region of radius 3 around the annotated cell centroid. **Prediction:** MS-D predicts probability maps of cell positions from the raw image. These centroid probabilities are thresholded and used to initialize the active contour segmentation that segments the cells from the raw images. The workflow is schematically depicted for the general 3D case in Fig. 2.

#### Cell Localization

For the 2D cell localization we used the MS-D architecture, with a *width* of 8 (multi-scale feature channels), a *depth* of 8 and a kernel size of 3×3 (see [11] for details). As the loss function we used the 1 – *F_β_* score on the binary pixel labels between reference and prediction. This score combines precision (p) and recall (c) as 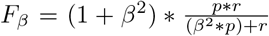. We set *β* = 0.7, putting more emphasize on precision of the predicted cell centroid regions. For optimization we used stochastic gradient descent with an adaptive learning rate (ADADELTA) [17]. We trained the network on randomly sampled image sections of size 256 × 256 pixels and the annotated centroids, convolved with a spherical kernel of radius 3. We used a batch size of 8 and applied data augmentation to the training samples in form of image rotation up to 90°. To derive cell centroids from the MS-D predictions, we thresholded the predictions at 70% probability level.

**Fig. 2.**
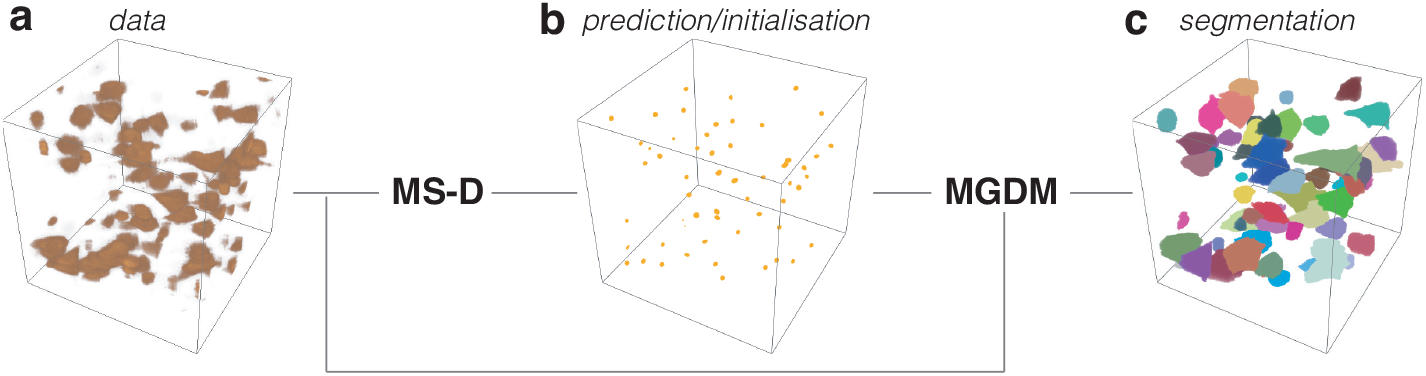
Schematic overview of method in 3D. We train a MS-D network on manually labeled cell centroids. The predicted cell positions are used as initialization and topology prior for the multi-object geometric deformable model (MGDM).

*Extension to 3D.* For the 3D images we used a 5×8 (width×depth) MS-D to fit into the GPU memory, with all 2D convolutional layers replaced by 3D convolutions with corresponding kernel size of 3×3×3.

We trained the network, with a batch size of 1 and without augmentation. As input we used pairs of randomly sampled image sections of size 96× 96× 96 pixels and the annotated centroids, convolved with a 3D spherical kernel of radius 3. Predictions were thresholded at 70% probability level.

#### Multi-object Geometric Deformable Model

Once cell centroids have been detected, the final segmentation is handled by a Multi-object Geometric Deformable Model (MGDM), an extension of the classical Deformable Geometric Model which ensures fast segmentation of an arbitrarily large number of cells while enforcing topological constraints between them [1]. The deformable model can be driven by any number of active contour forces. For simplicity, we only include balloon forces derived from the microscopy images and curvature regularisation, as follows.

For each detected cell *c*, we first find the maximum intensity *M_c_* of the microscopy image inside the non-zero probability region around each detected centroid. We set the MGDM balloon forces to decrease linearly with the distance to *M_c_*:

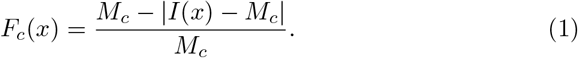

where *I*(*x*) is the image intensity. Because fluorescence intensity varies between cells, this calibration ensures that each cell is within its detection range. For the background *b*, we first estimate the mean overall image intensity *M_b_* (assuming that there is significantly more background than cells) to separate background from cells and derive a similar balloon force:

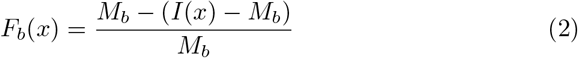

To avoid unstable evolution from too large forces, *F_c_* and *F_b_* are all bounded in [+1, —1]. These specific balloon forces are combined with classical curvature regularisation forces in the MGDM evolution equation, with *ϕ_c_, ϕ_b_* the signed distance function level sets defining the implicit curve evolution and *κ_c_, κ_b_* the corresponding level set mean curvature for cells *c* and background *b*:

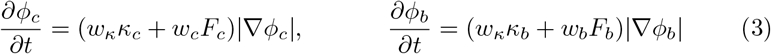

We used the same parameters for all 2D and 3D studies and fixed the weights for curvature regularisation to *w_κ_* = 0.6, and the balloon forces to *w_c_* = *w_b_* = 0.3. The evolution was run for 200 iterations.

**Fig. 3.**
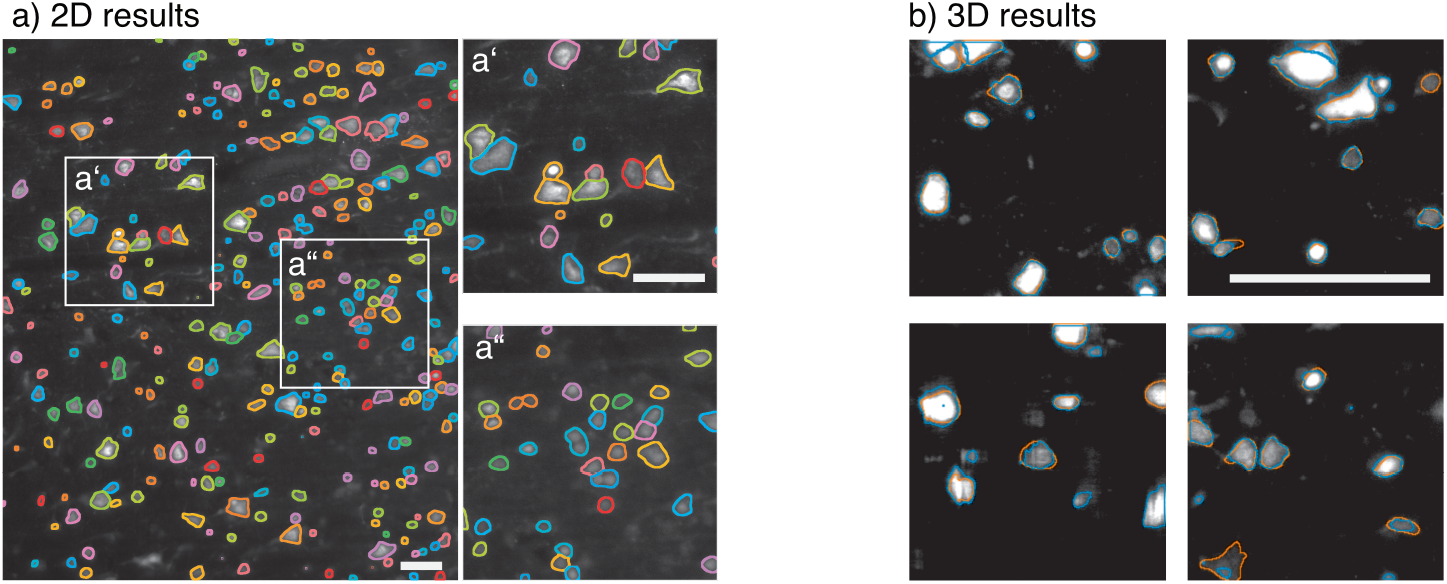
Segmentation result. (a) An image from the 2D test set showing (randomly colored) outlines of segmented cell masks, subregions were magnified for better visibility. (b) Example planes from the 3D test set showing the outlines of annotated reference masks (red) and segmentation results (blue). (All scale bars 100 *μm*.)

## 3 Results

### 2D cell localization and segmentation

To assess cell localization accuracy, we compared the MS-D net prediction to the manually annotated reference centroids and computed precision, recall, and the combined F1-score for the annotated images. Examples of final segmentation masks on the test set are shown in Fig.3a. The segmentation performed well across regions with varying cell appearance and density. The cell localization step produced a few fusion and splitting errors, particularly in regions where small, dim cells were concentrated (Fig.3a”). Quantitative results were aggregated over all four test images and summarized in Table 1. The proposed method improved cell detection and segmentation accuracy. We used the fastER segmentation [5] as a reference baseline because it produces state-of-the art results on par with deep learning methods such as [12]

**Table 1.** Comparison of segmentation across test set. Best results shown in bold.

| method | precision | recall | F1-score | JI(median) |
| --- | --- | --- | --- | --- |
| ours | **0.895** | **0.942** | **0.918** | **0.672** |
| adapt. threshold | 0.255 | 0.855 | 0.393 | 0.634 |
| FastER | 0.834 | 0.917 | 0.874 | 0.521 |

and can be trained with few annotations^7^. We further compare our results to a simple baseline using adaptive thresholding.

### 3D cell localization and segmentation

Examples of segmentation results on a test sample are shown in Fig.2c and on 2D sub-slices in Fig.3b (blue contours). Over the three test stacks our method achieved a good performance in 3D with a cell localization precision of 0.81, a recall of 0.873 resulting in an F1-score of 0.84 and a median Jaccard index of 0.732 for segmentation. The MGDM segmentation tended to result in slightly larger 3D masks compared to the manual reference as illustrated in Fig.3b showing the annotated outlines in red and the segmentation result in blue.

## 4 Conclusions

As a proof of concept we present a hybrid strategy to segment neural cells combining a mixed-scale neural network and a topology-preserving geometric deformable model. Our method robustly detects and segments cell bodies in light-sheet microscopy images of cleared post mortem human brain tissue. High quality results were obtained despite large variations in cell shape and intensity, anisotropic resolution and challenging imaging artifacts. Our method works for 2D and 3D images and only requires sparse centroid annotations for training. This is a crucial prerequisite for large-scale histological analysis of desired quality as fully annotated cell segmentations are very tedious and error-prone in 3D.

While there exist many different options for multi-label segmentation given the initial detected cell centroids, such as watersheds, graph cuts, or belief propagation, we chose multi-object geometric deformable models for their ability to constrain the cell boundary curvature and enforce topological relationships while allowing for a flexible design of the segmentation cost function based on local rather than global intensity variations. Thus, the method can be adapted to alternative clearing, staining and imaging protocols.

As a next step we will systematically optimize the neural network architecture, the loss function, and the forces of the MGDM segmentation to improve cell localization and segmentation further. Another interesting advance would be to train ensembles of networks to take the inter-network variability of predictions into account for downstream processing.

## Footnotes

6 Related ideas integrating deep learning and level set formulations have been proposed by [14] or [6]. In contrast to our approach based on sparse centroid annotations these methods require pixel-accurate object masks for training.

7 Note that fastER is limited to 2D images only.

